# Sequence features around cleavage sites are highly conserved among different species and a critical determinant for RNA cleavage position across eukaryotes

**DOI:** 10.1101/2021.03.10.434873

**Authors:** Daishin Ueno, Shotaro Yamasaki, Yuta Sadakiyo, Takumi Teruyama, Taku Demura, Ko Kato

**Affiliations:** Graduate School of Science and Technology, Nara Institute of Science and Technology, Ikoma, 630-0192 Japan

**Keywords:** RNA degradation, degradome sequencing, machine learning

## Abstract

RNA degradation is critical for control of gene expression, and endonucleolytic cleavage– dependent RNA degradation is conserved among eukaryotes. Some cleavage sites are secondarily capped in the cytoplasm and identified using the CAGE method. Although uncapped cleavage sites are widespread in eukaryotes, comparatively little information has been obtained about these sites using CAGE-based degradome analysis. Previously, we developed the truncated RNA-end sequencing (TREseq) method in plant species and used it to acquire comprehensive information about uncapped cleavage sites; we observed G-rich sequences near cleavage sites. However, it remains unclear whether this finding is general to other eukaryotes. In this study, we conducted TREseq analyses in fruit flies (*Drosophila melanogaster*) and budding yeast (*Saccharomyces cerevisiae*). The results revealed specific sequence features related to RNA cleavage in *D. melanogaster* and *S. cerevisiae* that were similar to sequence patterns in *Arabidopsis thaliana*. Although previous studies suggest that ribosome movements are important for determining cleavage position, feature selection using a random forest classifier showed that sequences around cleavage sites were major determinant for cleaved or uncleaved sites. Together, our results suggest that sequence features around cleavage sites are critical for determining cleavage position, and that sequence-specific endonucleolytic cleavage–dependent RNA degradation is highly conserved across eukaryotes.

## INTRODUCTION

Multiple mechanisms are involved in control of gene expression. RNA degradation is a critical step in maintaining RNA homeostasis in eukaryotes (Keene 2010). RNA degradation is initiated either by deadenylation followed by removal of the cap and 5’-to-3’ or 3’-to-5’ exonucleolytic digestion, or by endonucleolytic cleavage followed by exonucleolytic digestion (Parker 2012). Deadenylation-dependent RNA degradation is well studied in yeast, plants, and animals, and proteins and sequences related to deadenylation or decapping have been characterized in detail (Wu and Brewer 2008; Chiba and Green 2009; Parker 2012). Although microRNA- and siRNA-dependent forms of RNA degradation are well studied in eukaryotes, endonucleolytic cleavage–dependent RNA degradation has not been analyzed in detail.

Previous studies developed several degradome sequencing methods to detect cleavage sites in plants such as *A. thaliana*, mainly in the context of micro RNA- or siRNA-mediated cleavage (Addo-Quaye et al. 2008; German et al. 2008; Gregory et al. 2008). Using these methods, RNA cleavage sites were identified in plants (Zhou et al. 2010; Shamimuzzaman and Vodkin 2012; Karlova et al. 2013; Hou et al. 2016; Anderson et al. 2018), yeast, and animal models (Harigaya and Parker 2012; Bracken et al. 2011). However, because these methods selected poly(A) RNA (Willmann et al. 2014; Addo-Quaye et al. 2008), many of the 5’ degradation intermediates they detected appeared to be biased toward the 3’-ends of transcripts (Weinberg et al. 2016). In addition, ligation of single-stranded 5’ adapters using T4 RNA ligase induces bias in the detected 5’ RNA ends (Hou et al. 2014; Zhuang et al. 2012). Specifically, ligation efficiencies are quite low if there are stable secondary structures between adapters and 5’-proximal RNA sequences (e.g., high G/ C frequency). In addition, non-specific annealing occurs when using single-stranded 5’ adapters and subsequent PCR amplification, resulting in sequence artifacts (Hou et al. 2014).

Alternatively, instead of previous degradome methods, Cap analysis of gene expression (CAGE) can be used to detect cleavage sites (Mercer et al. 2010). To decrease the bias in 5’ adapter ligation efficiency, the CAGE method joins single-strand cDNAs to partially double-stranded adapter sequences in the presence of polyethylene glycol (PEG) (Murata et al. 2014). The partially double-stranded adapter sequence has phosphate modifications, allowing a dramatic reduction in 5’ adapter dimers while simultaneously increasing adapter ligation efficiency (Takahashi et al. 2012). In addition, formation of secondary structure between adapter and 5’-proximal RNA sequences is less frequent for the partially double-stranded adapter sequences than for single-stranded adapter sequences, and ligation efficiencies are significantly increased by the addition of PEG to the ligation buffer (Song et al. 2014). Furthermore, current CAGE methods do not involve PCR amplification or restriction enzyme digestion (Murata et al. 2014), allowing CAGE to detect 5’ RNA ends more accurately than other methods (Adiconis et al. 2018). However, because the CAGE method was developed for detecting transcription start sites (TSSs), the majority of 5’ RNA ends are TSSs, and expected cleavage sites constituted less than 20% of all detected sites (Mercer et al. 2010). Also, because some cleavage sites are secondarily capped in the cytoplasm (Schoenberg and Maquat 2009) and the CAGE method selects Cap RNA using Cap-trapping methods, the detected cleavage sites are likely to be re-capped. Uncapped RNAs are prevalent in eukaryotes (Ibrahim et al. 2018; Bracken et al. 2011) but the majority of uncapped cleavage sites cannot be detected using CAGE methods.

In a previous study, we improved these features by developing truncated RNA-end sequencing (TREseq) based on the no-amplification non-tagging CAGE (n An T-i CAGE) method in *A. thaliana* (Ueno et al. 2018). Initially, we separated RNA into Cap and Cap-less RNA using Cap-t rapping method and constructed libraries for Cap-less RNA, in addition to Cap RNA. To increase the concentration of Cap-less RNA, we removed the majority of expected Cap RNA from the Cap-less RNA data during the mapping process. Because Cap (7-methylguanosine) is not encoded in the genome and nucleotide of Cap position in the genome is Y (C or T) in eukaryotes (Carninci et al. 2006; Adiconis et al. 2018; Thieffry et al. 2020; Lu and Lin 2019), the majority of Cap RNA in the Cap-less data could be removed by excluding Cap-less RNA with mismatches at the 5’ end of the read relative to the genome (Ueno et al. 2020). In addition, we decreased the 3’ bias of the cleavage site using random primers and conducting rRNA depletion (Ueno et al. 2018). By improving these experimental procedures, we acquired comprehensive information about uncapped cleavage sites and identified a relationship between cleavage efficiencies and RNA stability. Although previous degradome methods cannot identify such relationships, we observed that genes with high cleavage efficiencies had short half-lives relative to those with lower cleavage efficiencies (Ueno et al. 2018). When we focused on nucleotide f requencies, we observed G-rich sequence motifs around the cleavage sites; moreover, genes with G-rich sequence motifs accumulated lower RNA levels than genes without motifs (Ueno et al. 2020). These tendencies were conserved not only in *A. thaliana* but also among different plant species (Ueno et al. 2018, 2020). In addition, cleavage sites were common around the start and stop codon, and three-nucleotide periodicity was observed in coding region sequences (CDSs) in plant species (Ueno et al. 2020). These tendencies are similar to ribosome movements, which have been reported by ribosome profiling methods, suggesting that the translation process is related to RNA cleavage and that ribosome position is important for determining the position of cleavage sites (5’ truncated RNA ends) (Ibrahim et al. 2018; Yu et al. 2016; Pelechano et al. 2015). Although specific sequence features around cleavage sites are observed in plant species, it remains to be determined whether these properties are shared with other eukaryotes. In addition, RNA cleavage position seems to be determined by multiple determinants (e.g., sequence features and translation), but previous studies have not performed an integrated analysis of these determinants.

To address this issue, we conducted TREseq in fruit flies (*Drosophila melanogaster*) and budding yeast (*Saccharomyces cerevisiae*), and obtained TREseq data in *A. thaliana* from a previous study. We also obtained ribosome profiling data and analyzed the effects of the translation process and sequence features on RNA cleavage across species. In addition, we conducted feature selection using a random forest classifier and evaluated the significance of the effects of multiple determinants on RNA cleavage positions in various species. Our research revealed that G-nucleotide frequencies were higher around cleavage sites in *D. melanogaster* and *S. cerevisiae*, similar to sequence features around cleavage sites in *A. thaliana,* and that these sequence features contribute to control of RNA cleavage across eukaryotes.

## RESULTS

### Improvement of experiment procedures and validation of cleavage sites in TREseq

We obtained TREseq data of *D. melanogaster* and *S. cerevisiae* in this study and *A. thaliana* (DRA 005995) from previous study (Ueno et al. 2018) (Supplementary Figure S1). In a previous study, we developed the TREseq method to decrease the bias towards the 3’ end of the transcript (Ueno et al. 2018). TREseq can detect cleavage sites both with [poly (A) ^+^] and without poly A tails [poly (A)^-^] using random primers, decreasing the apparent bias. In this analysis, the cleavage sites were almost equally distributed across genes in *D. melanogaster* and *S. cerevisiae*, as in *A. thaliana* (Supplementary Figure S2). If this improvement of the bias towards the 3’ end of the RNA was caused by selecting poly (A) ^-^ cleavage sites in addition to poly (A) ^+^ cleavage sites, we should observe similar tendencies when poly (A) ^-^ cleavage sites are isolated during library preparation. In the parallel analysis of RNA ends (PARE) method, poly (A)^-^ cleavage sites are selected, and poly (A) tail are added to the 3’ ends of cleavage sites before reverse transcription using oligo dT primer (Nagarajan et al. 2019) (Supplementary Figure 3). Using data obtained by the PARE method in *A. thaliana*, we compared the distribution patterns of cleavage sites in libraries containing RNA with and without poly(A) tails. The 3’ bias of cleavage sites was decreased by selection of poly (A) ^-^ cleavage sites, even when oligo dT primer was used in reverse transcription. These results strongly suggest that the bias of detected cleavage sites toward the 3’ end of transcripts was the result of selecting poly (A) ^+^ cleavage sites during library preparation.

In previous study, cleavage sites were detected using CAGE, but a low concentration of Cap-less RNA prevented high coverage (read depth) of uncapped cleavage site data. Therefore, we sought to determine how much TREseq could increase the concentration of Cap-less RNA. We added synthetic Cap-less *R-luc* RNA to total RNA derived from *A. thaliana* and separated RNA into Cap RNA or Cap-less RNA according to the method for TREseq library preparation (Supplementary Figure S4A). Subsequently, we quantified the abundance of Cap-less *R-luc* RNA in the Cap RNA and Cap-less RNA libraries using q RT-PCR. Relative to the Cap RNA library, Cap-less *Rluc* RNA was 10-fold more abundant in the Cap-less RNA library (Supplementary Figure S4B). We also confirmed that RNA could be clearly separated into Cap and Cap-less RNA by comparing a TREseq analysis in *A. thaliana* that did not use the Cap-trapping method for library preparation (Supplementary Figure S4C). The results revealed that Cap-less RNA was highly concentrated using the Cap-trapping method, and that the degree of Cap RNA contamination in Cap-less RNA data was low (Supplementary Figure S4C). The same is true for Cap R NA data (Supplementary Figure S4C). These results indicate that TREseq yields a higher concentration of Cap-less RNA than the CAGE method.

To detect 5’ RNA ends, several methods were developed in previous studies, in particular for detection of transcription start sites (Adiconis et al. 2018). We can roughly classify methods into several groups: ligation of partially double-stranded adapter (CAGE or TREseq), template switching (RAMPAGE, STRT, and NanoCAGE XL), and ligation of single-stranded adapter (previous degradome analysis) (Adiconis et al. 2018). Although nanoCAGE (template switching), which was developed for detecting TSSs in nanogram ranges of total RNA, included sequence artifacts, the experimental procedures were improved in the present CAGE method, which can detect 5’ RNA ends more accurately than the other methods (Adiconis et al. 2018; Murata et al. 2014). Because TREseq was developed based on CAGE (n An T-iCAGE) and uses partially double-strand adapter sequences (no PCR amplification), sequence artifacts reported in previous degradome methods (Hou et al. 2014) theoretically should not occur. Although expected cleavage sites detected by CAGE are validated by 5’ RACE (Carninci et al. 2006), and cleavage positions in microRNA, siRNA, or long non-coding RNAs (lncRNAs) detected by CAGE matched those in the database (Mercer et al. 2010; Nepal et al. 2013; De Rie et al. 2017), we also confirmed that cleavage sites in TREseq could be detected using different methods in *A. thaliana*. As a different method, we used Nanopore sequencing (template-switching method). We selected 3000 cleavage sites whose detected reads were highest in each gene, and found that these sites were associated with high G-nucleotide frequencies (Supplementary Figure S5A), similar to the sequence features around sites with high cleavage efficiencies identified by TREseq in *A. thaliana* (Ueno et al. 2018). A Nanopore sequencer can detect 5’ RNA ends similarly to a short-read sequencer (Parker et al. 2020), but accurate mapping in continuous identical nucleotides (e.g., G-rich sequences) is more difficult at one-nucleotide resolution. Although the peaks were slightly shifted, we observed similar tendencies between TREseq and Nanopore sequencing (Supplementary Figure S5B). These results suggest that the detected cleavage sites are reliable, and that high G-nucleotide frequencies around cleavage sites are not sequence artifacts of TREseq.

Based on these results, we concluded that TREseq can decrease the bias of detected cleavage sites toward the 3’ end of transcripts and improve the concentration of Cap-less RNA. In addition, we validated these sites by a different method (Nanopore sequencer), and suggesting that the cleavage sites detected by the TREseq method are reliable.

### Calculating the cleavage score (CS) in TREseq

In degradome analysis, the term “cleavage score” was used to estimate RNA cleavage efficiencies in previous studies (Anderson et al. 2018; Ueno et al. 2018). In this analysis, we defined cleavage score at the site level (CS_site_) as the number of 5’ degradation intermediates normalized against RNA abundance (Ueno et al. 2018, 2020). In addition, as with efficiencies of RNA cleavage at the gene level, we defined the sum of CS_site_ values in each gene as the CS_gene_ value. We calculated CS_site_ and CS_gene_ values in *S. cerevisiae*, *D. melanogaster* and *A. thaliana* (Supplementary Figures S6-8). We confirmed the reproducibility of CS_site_ values in two biological replicates (Supplementary Figure S9); the CS_site_ values were almost normally distributed in both species, as in *A. thaliana* (Supplementary Figure S9).

### Cleavage-dependent degradation mechanisms are important for controlling RNA stability across species

Initially, we compared the CS_gene_ values with RNA half-lives obtained from previous studies (Narsai et al. 2007; Neymotin et al. 2014; Burow et al. 2018). CS_gene_ values were related to half-life information in a previous study (Ueno et al. 2018), and we confirmed that genes with high CS_gene_ values had shorter half-lives using different half-life measurement data from *A. thaliana* (Figure 1). CS_gene_ values and half-lives were negatively correlated in *D. melanogaster* and *S.* cerevisiae, as well as in *A. thaliana* (Figure 1). When we selected genes with TOP and BOTTOM 10% CS_gene_ values, similar tendencies were observed in *A. thaliana* (median = 3.4 h in TOP 10%; median = 8.1 h in BOTTOM 10%), *D. melanogaster* (median = 74 min in TOP 10%; median = 110 min in BOTTOM 10%) and *S. cerevisiae* (median = 7.5 min in TOP 10%; median = 27 min in BOTTOM 10%). Gene ontology enrichment analysis revealed that regulation of transcription, phosphorylation, signal transduction, and cell cycle processes were enriched in genes with high CS_gene_ values, and that translation was enriched in genes with low CS_gene_ values, across species (Supplementary Tables S1, S2). These results are consistent with previous RNA half-life analyses concluding that RNAs with short half-lives were quick responses to stimuli, whereas RNAs with long half-lives tended to be involved in processes related to constant maintenance of cellular physiology (Tani et al. 2012; Narsai et al. 2007; Ueno et al. 2018). Although the previous method could not identify the relationship between cleavage efficiencies and R NA stability, we found that genes with high CS_gene_ values have short half-lives. These results were obtained due to our improvement of the experimental procedures. Together, these results indicate that cleavage-dependent RNA degradation is important for controlling RNA stability and contributes to various biological processes across eukaryotes.

**Figure 1.**
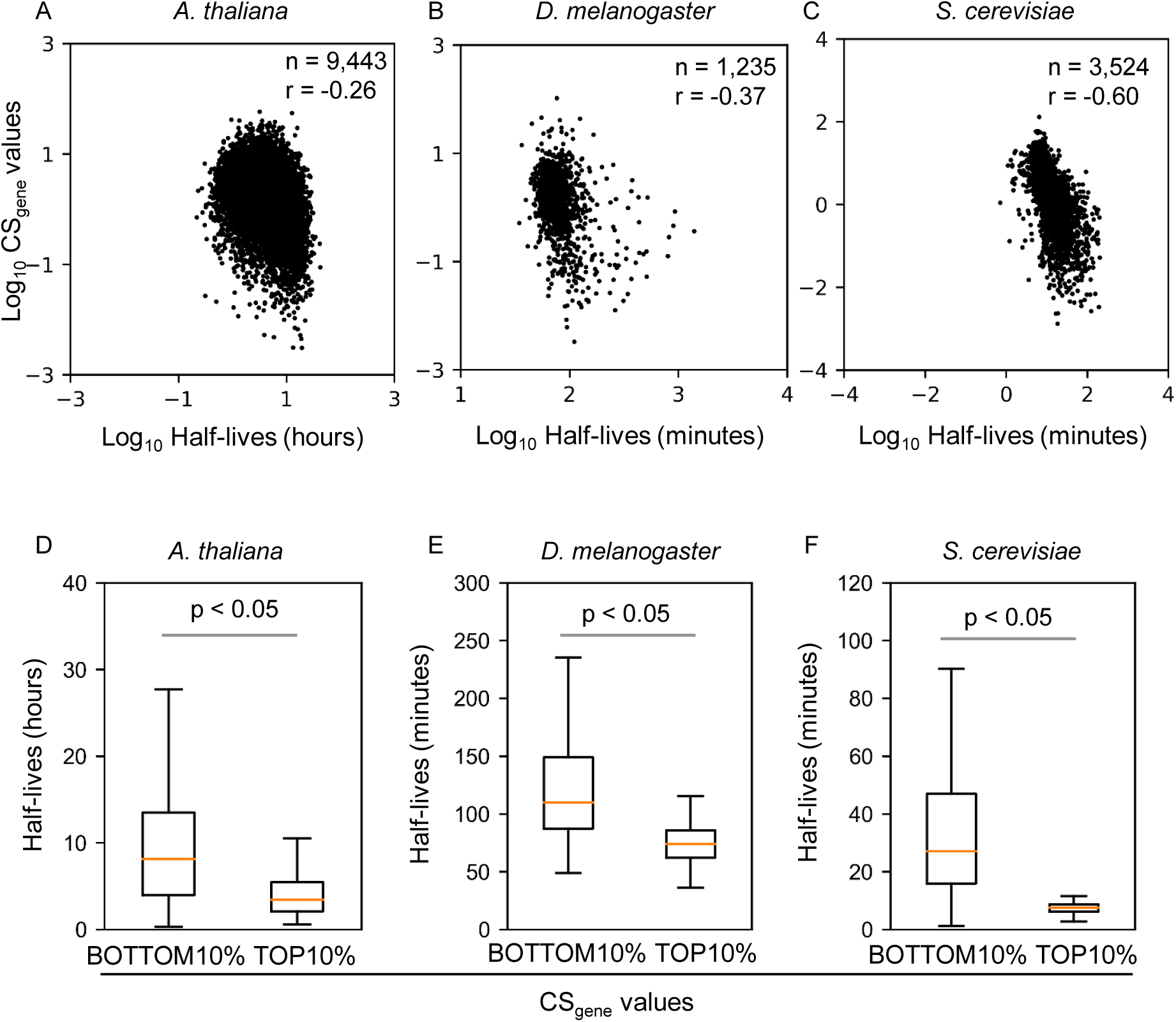
Relationship between CS_gene_ values and RNA half-lives. Scatter plots of half-life vs. CS_gene_ values in *A. thaliana* (A), *D. melanogaster* (B), and *S. cerevisiae* (C). We selected the TOP and BOTTOM 10% of CS_gene_ values and compared half-lives in each species (D– F). Welch’s t-test (two-sided) was used to determine statistical significance. The number of genes analyzed in the TOP 10% and BOTTOM 10% were 944 genes in *A. thaliana*, 124 in *D. melanogaster*, and 352 in *S. cerevisiae*. Outliers were removed from box plots.

### Conservation of sequence features around cleavage sites

Less information about untranslated regions (UTRs) is available in *D. melanogaster* and *S. cerevisiae* than in *A. thaliana*. Therefore, in our subsequent analysis, we compared the sequence features of cleavage sites using CDS regions. When we focused on nucleotide frequencies around the cleavage sites, we observed similar sequence features among species (Figure 2). In addition, sequence features around cleavage sites could be more clearly observed for sequences with high cleavage efficiencies in *D. melanogaster* and *S. cerevisiae*, as in *A. thaliana* (Figure 2).

**Figure 2.**
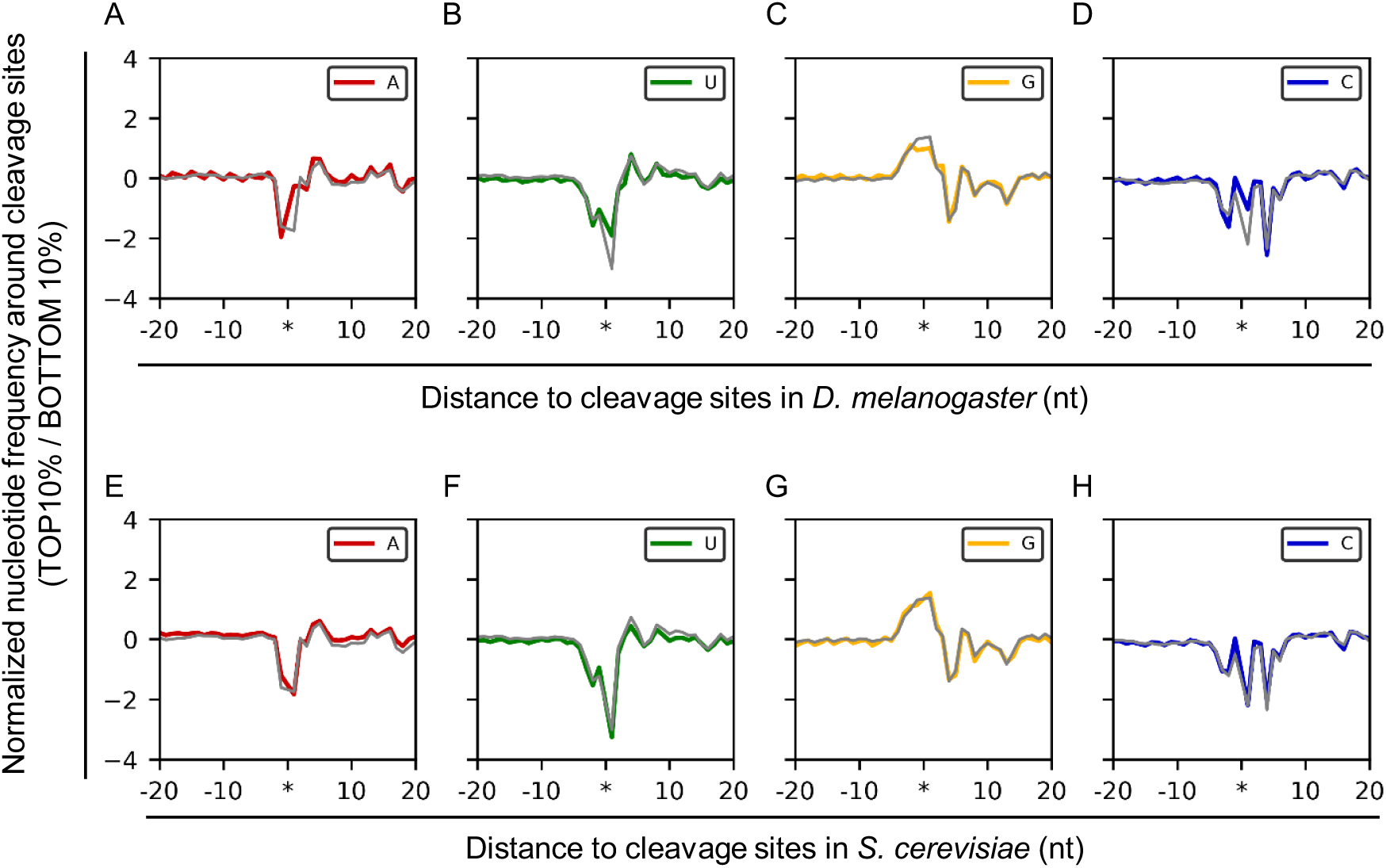
Nucleotide frequency around the cleavage sites. We selected the TOP and BOTTOM 10% cleavage sites based on CS_site_ values and calculated normalized nucleotide frequency (log_2_ ratio of nucleotide frequencies between TOP and BOTTOM 10%) in *D. melanogaster* (60,096 sites analyzed in TOP and BOTTOM 10%) and *S. cerevisiae* (61,689 sites analyzed in TOP and BOTTOM 10%). We compared normalized nucleotide frequency around cleavage sites between *A. thaliana* and *D. melanogaster* (A–D) or *S. cerevisiae* (E–H). Gray line indicates nucleotide frequency in *A. thaliana*. Asterisks indicate positions betwee n-1 and +1.

Subsequently, we discovered sequence motifs using sequences around (from − 10 to + 10), upstream (from − 20 to − 5), or downstream (from + 5 to + 20) of cleavage sites using DREME, and then used STAMP to carry out phylogenetic analyses based on similarities among the discovered motifs (Supplementary Figures S10–S12). We also obtained sequences around, upstream, or downstream of cleavage sites in different plant species (*Lactuca sativa* [lettuce], *Oryza sativa* [rice], and *Rosa hybrida* [rose]) reported in a previous study (Ueno et al. 2020), and added sequence motifs for phylogenetic analyses. Generally, similar G-rich sequence motifs were observed around the cleavage sites (from − 10 to + 10). By contrast, A- or AU-rich motifs were observed upstream or downstream of cleavage sites in plants or yeast, and different sequence motifs were observed in *D. melanogaster*. These results were consistent with sequence conservation in RNA 3’-processing sites, indicating that similar sequences were observed around processing sites, but sequences upstream or downstream of processing sites were not shared among species (Tian and Graber 2012). Specifically, sequences were quite similar between yeast and plants, but differed in animals (Tian and Graber 2012). These results suggest that similar sequence features around cleavage sites are conserved across eukaryotes, and that different sequences in upstream or downstream regions, in addition to highly conserved sequence around cleavage sites, also contribute to control of cleavage efficiencies in different species.

Because G-rich sequences sometimes induce stable RNA structures, which play important roles in biological process (Lipps and Rhodes 2009; Subramanian et al. 2011; Millevoi et al. 2012), we subsequently focused on the RNA structures around cleavage sites. We predicted paired or unpaired site (dot-bracket notation) around cleavage sites using RNAfold and base-paring frequencies at each position were calculated in each species. We observed that base-pairing frequency in TOP 10% CS_site_ values were higher around cleavage sites compared to BOTTOM 10% CS_site_ values in all species (Supplementary Figure S13A–C). In addition, pairing-frequency was dependent on G-nucleotide frequencies around the cleavage sites (Supplementary Figure S13D–E). These results suggest that RNA structure is related to RNA cleavage, and that these structures are dependent on G-nucleotide frequency around cleavage sites.

In our analysis, we observed specific sequence features around cleavage sites and hypothesized that these sequences were important for determining cleavage position. We extracted sequences from − 5 to + 15 nucleotides around the cleavage sites, and then created sequence motifs (motif letter-probability matrix lines) based on frequencies. Subsequently, we calculated the distribution of sequence motifs in each gene using FIMO and compared it with the distribution of cleavage sites (Figure 3A). If sequence features are important for determining the cleavage position, we expected to observe the same distribution patterns between sequence motifs and cleavage sites. We found that sequence motif had similar three-nucleotide periodicities, and that the phases were almost same (Figure 3B–G). Although previous studies suggested that the three-nucleotide frequency is caused by ribosome movement (Pelechano et al. 2015; Ibrahim et a l. 2018; Yu et al. 2016), these results suggest that sequence features are also related to the three-nucleotide periodicity of cleavage sites and are important for determining RNA cleavage position.

**Figure 3.**
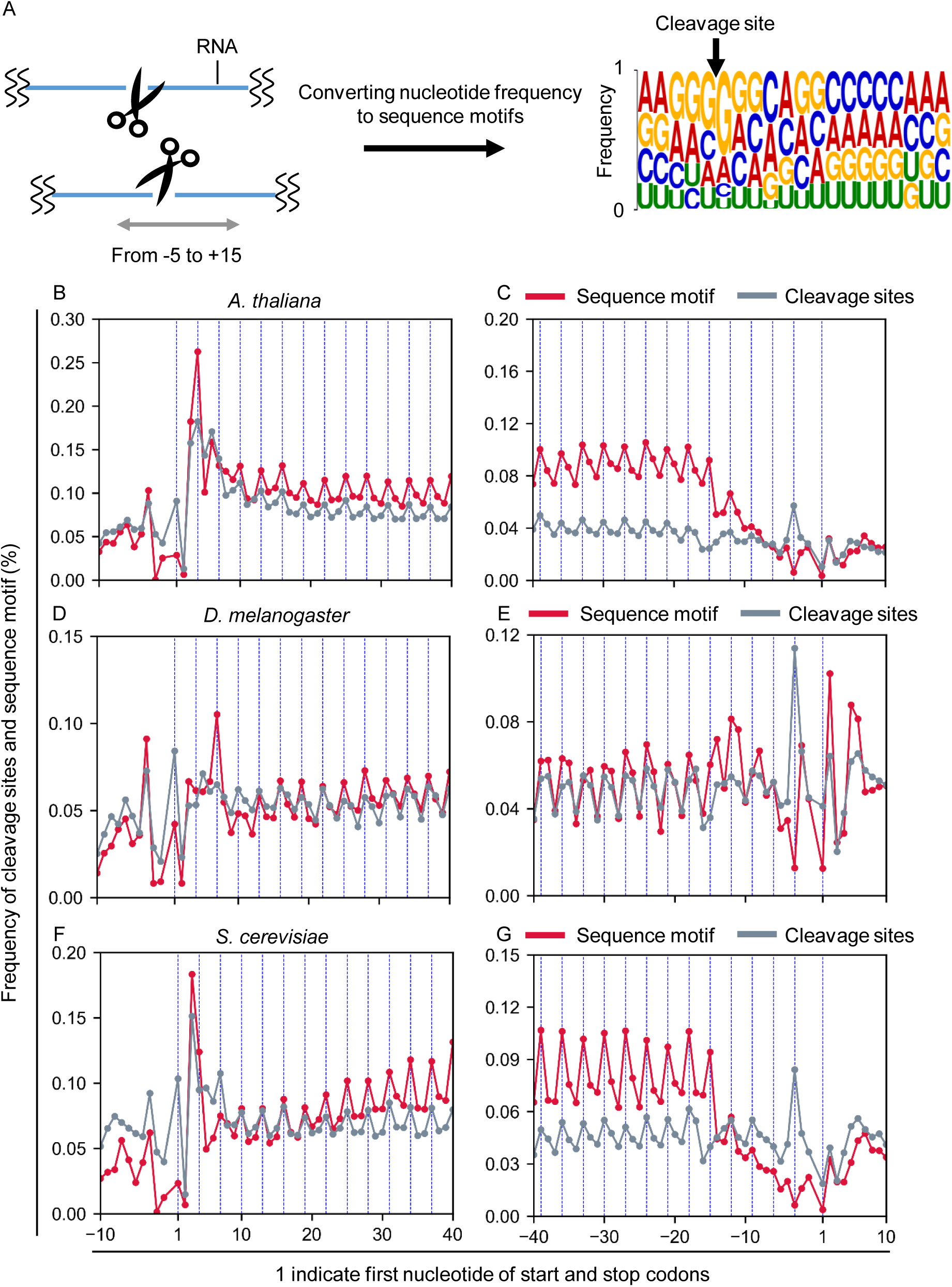
Distribution of cleavage sites and sequence motifs around cleaved sites. The nucleotide frequency information was converted into sequence motifs (motif letter-probability matrix lines) in MEME (A), and distribution of sequence motif was computed using FIMO. We compared the distribution pattern of cleavage sites and sequence motifs around start (B, D, F) and stop (C, E, G) codons in *A. thaliana* (B, C), *D. melanogaster* (D, E), and *S. cerevisiae* (F, G). Because most UTR information in *S. cerevisiae* is not yet defined, we expanded the annotation region of *S. cerevisiae* only in this analysis.

Together, these results indicate that sequence features around cleavage sites are highly conserved across species and are important for determining both cleavage efficiencies and cleavage positions in eukaryotes.

### Effects of the translation process on cleavage position and cleavage efficiency

Previous studies in yeast reported that changes in ribosome occupancy and degradation efficiencies at the gene level were positively correlated between oxidative stress and normal conditions (Pelechano et al. 2015). Hence, we analyzed the relationship between cleavage efficiencies and ribosome occupancy at the gene level among species. In addition, we analyzed the effect of ribosome occupancy on cleavage efficiency at the site level. For *A. thaliana*, we conducted ribosome profiling using the same conditions as for our previous TREseq data. For *D. melanogaster* and *S. cerevisiae*, we obtained ribosome profiling data from previous studies (Luo et al. 2018; Gerashchenko and Gladyshev 2017). We defined the number of RPFs at each site, normalized against RNA abundance, as the RO_site_ values, and the sum of RO_site_ values at the CDS regions of each gene as the RO_CDS_ values. Similarly, we defined the sum of CS_site_ values in the CDS region as the CS_CDS_ values. At the gene level, we observed a positive correlation between CS_CDS_ and RO_CDS_ values among species (Figure 4). These results suggest that the translation process has a positive effect on cleavage efficiencies at the gene level.

**Figure 4.**
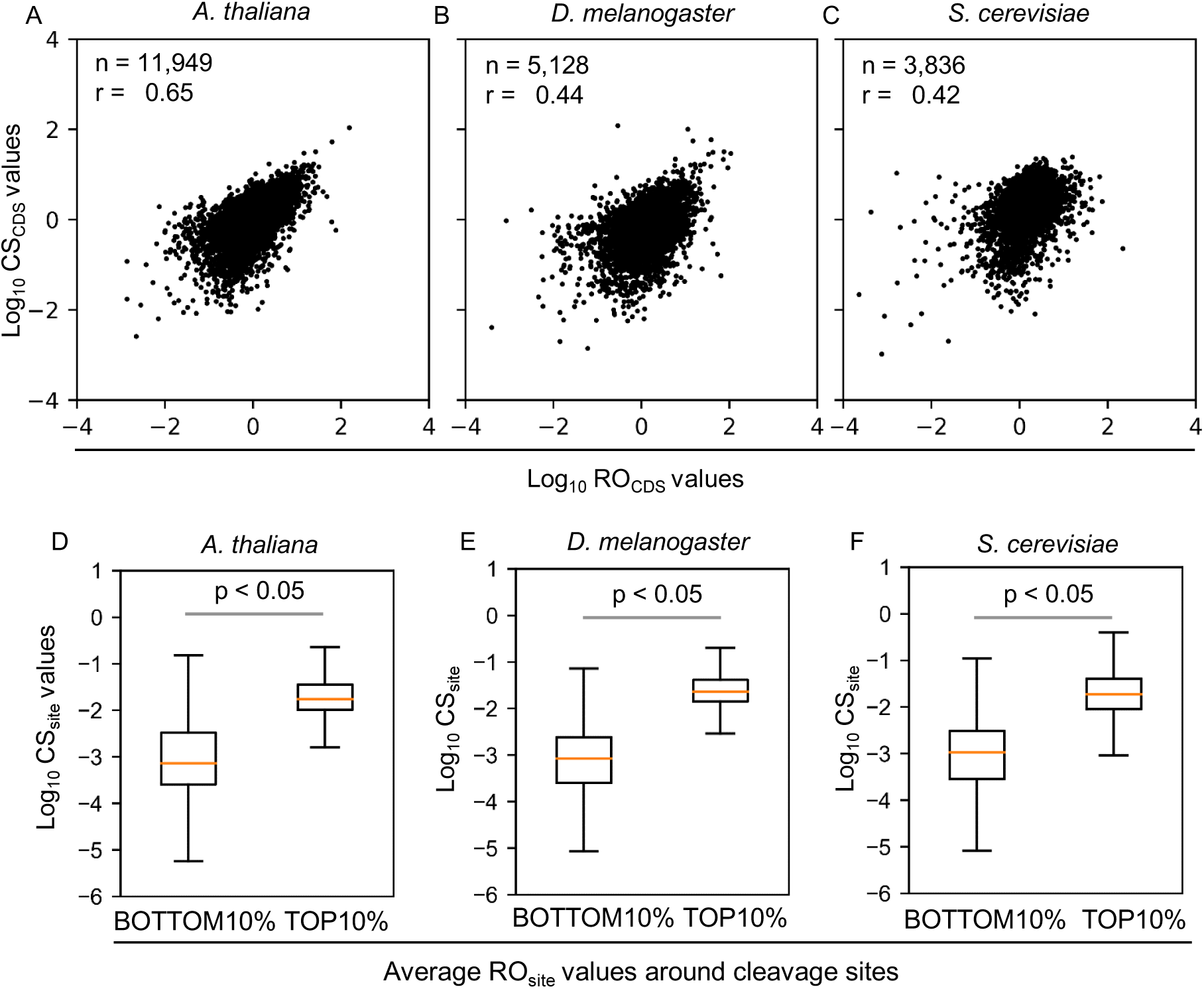
Relationship between ribosome occupancy and cleavage efficiencies. Scatter plots of CS_CDS_ and RO_CDS_ values in *A. thaliana* (A), *D. melanogaster* (B), and *S. cerevisiae* (C). Both CS_CDS_ and RO_CDS_ values were normalized against RNA length. Average ribosome occupancy around cleavage sites in the CDS region was calculated and compared against CS_site_ values in the TOP 10% (high ribosome occupancy) and BOTTOM 10% (low ribosome occupancy) (D–F). Welch’s t-test (two-sided) was used to determine statistical significance. The number of genes analyzed in the TOP 10% and BOTTOM 10% were 177,248 in *A. thaliana*, 60,096 in *D. melanogaster*, and 61,689 in *S. cerevisiae*. Outliers were removed from box plots.

Next, we focused on the relationship between CS_site_ values and ribosome occupancies at the site level. We calculated the average RO_site_ value around the cleavage sites (±50 nucleotides), and selected the TOP or BOTTOM 10% based on occupancy around the cleavage site (Figure 4). Even at the site level, cleavage sites with high ribosome occupancy (TOP 10%) had higher cleavage efficiencies than the BOTTOM 10% across species. These results suggested that ribosome occupancy has a positive effect on cleavage efficiencies at the gene or site level. Although correlations between ribosome occupancy and degradation efficiency were weaker under normal conditions compared to correlations between changes in ribosome occupancy and degradation efficiency between under normal condition and oxidative stress in yeast (Pelechano et al. 2015), we could observe a positive correlation among all species examined. Because the previous study conducted poly(A) selection, resulting in a bias in detection level dependent on RNA length (Weinberg et al. 2016), the correlation appeared to be lower than that detected by TREseq analysis.

Because previous study suggested that ribosome movement is related to cleavage position (5’-truncated RNA ends) (Pelechano et al. 2015; Yu et al. 2016; Ibrahim et al. 2018), we also compared the distributions of ribosomes and cleavage positions (Figure 5). We observed a three-nucleotide periodicity and found that both patterns were matched in some regions. However, the phases were generally not matched between cleavage and ribosome positions across different species. These results were consistent with a previous study showing that ribosome and cleavage positions (5’-truncated RNA ends) have three-nucleotide periodicities, but the phases are generally not matched (Ibrahim et al. 2018; Pelechano et al. 2015). These results suggest that ribosome position has effects on cleavage position (5’-truncated RNA ends) at a certain distance (Pelechano et al. 2015) or is not a major determinant for cleavage position. Together, these findings indicate that ribosome occupancies are important for cleavage efficiencies, but the effect of ribosome movement on cleavage position cannot be clarified solely by comparing the distribution patterns of ribosome and cleavage position.

**Figure 5.**
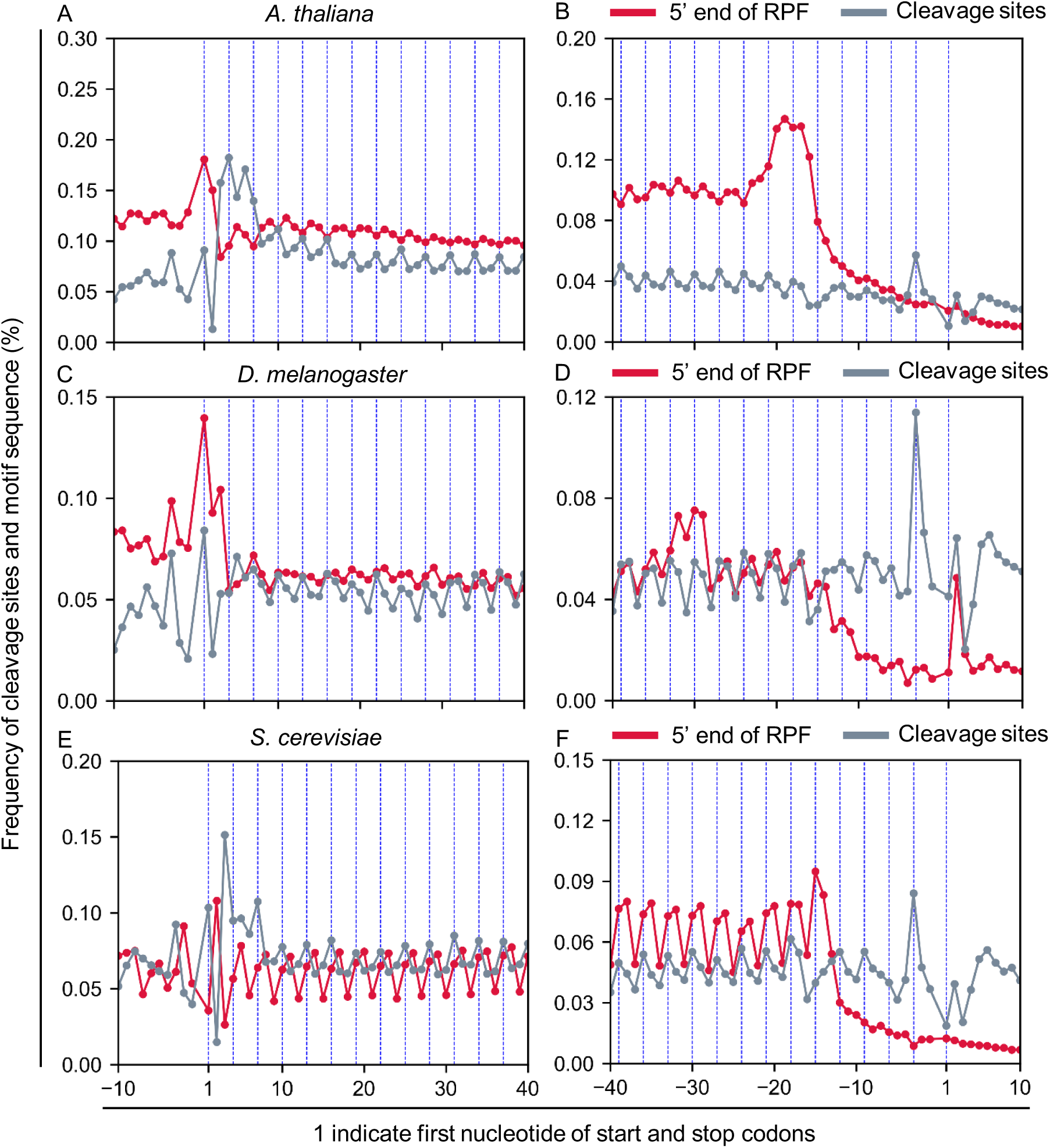
Comparing distributions of ribosome and cleavage positions. Distributions of 5’ ends of RPFs (ribosome protected fragments) and cleavage sites around start (A, C, E) and stop (B, D, F) codons in *A. thaliana* (A, B), *D. melanogaster* (C, D), and *S. cerevisiae* (E, F). Because most UTR information in *S. cerevisiae* is not yet defined, we expanded the annotation region of *S. cerevisiae* only in this analysis.

### Feature selection for determining cleavage position

Previous studies suggest that ribosome position is important for determination of 5’-truncated RNA ends (cleavage position) (Pelechano et al. 2015; Yu et al. 2016; Ibrahim et al. 2018). However, less information is available about the relationship between ribosome and cleavage positions, and we did not detect a strong effect of ribosome movement on cleavage position in this analysis. To evaluate the influence of sequences or ribosome movement on cleavage position, we conducted feature selection using a random forest (RF) classifier, which allows us to evaluate several features (determinants) according to the coefficient in the model. We selected sites and predicted whether they were cleaved (Figure 6A). To simplify the interpretation, we selected the site with the highest CS_site_ value in each gene, as well as sites with no CS_site_ values. For our model, we used nucleotide frequency, secondary structure, and ribosome position or ribosome occupancy information (Supplementary Figure S14). Ribosome position indicates only positional information, whereas ribosome occupancy includes occupancy and positional information. To evaluate the effect of secondary structure (base-paring information), we predicted paired or unpaired sites (dot-bracket notation) around cleavage sites using RNAfold. We divided the data into training and test sets. The model was constructed using the training data, and model performance was evaluated using the test data (Supplementary Figure S15). To estimate prediction accuracy in the test data, we used ROC curve and AUC values (Singh et al. 2017; Lloréns-Rico et al. 2015). ROC curves indicate the true-positive and false-positive rates, whereas AUC values indicate the area under the ROC curve. AUC values can range from 0.5 (i. e., the method does not perform better than random) and 1 (the method classifies all samples perfectly with no misprediction) (Lloréns-Rico et al. 2015). Initially, we focused on *A. thaliana* and made four models: [1] all features, [2] only nucleotide frequency around cleavage sites, [3] only secondary structure around cleavage sites, and [4] only ribosome position and ribosome occupancy around the cleavage sites. AUC value (prediction accuracy) was highest in classification model that used all features (Figure 6B). In addition, when we used only nucleotide frequencies, the prediction accuracy was still high. By contrast, when we used only secondary structures or ribosome position/ occupancy information, prediction accuracy was lower. These results were consistent with the Gini importance, which indicate the importance of features for predicting whether site is cleaved or not, in RF classifier using all features (Supplementary Figure S16). These tendencies were also observed in *D. melanogaster* or *S. cerevisiae* (Supplementary Figures S17–S19). Although ribosome movement also seemed to be related to determination of cleavage position based on the Gini importance, these results suggest that sequence features are the major determinant of cleavage position across eukaryotes.

**Figure 6.**
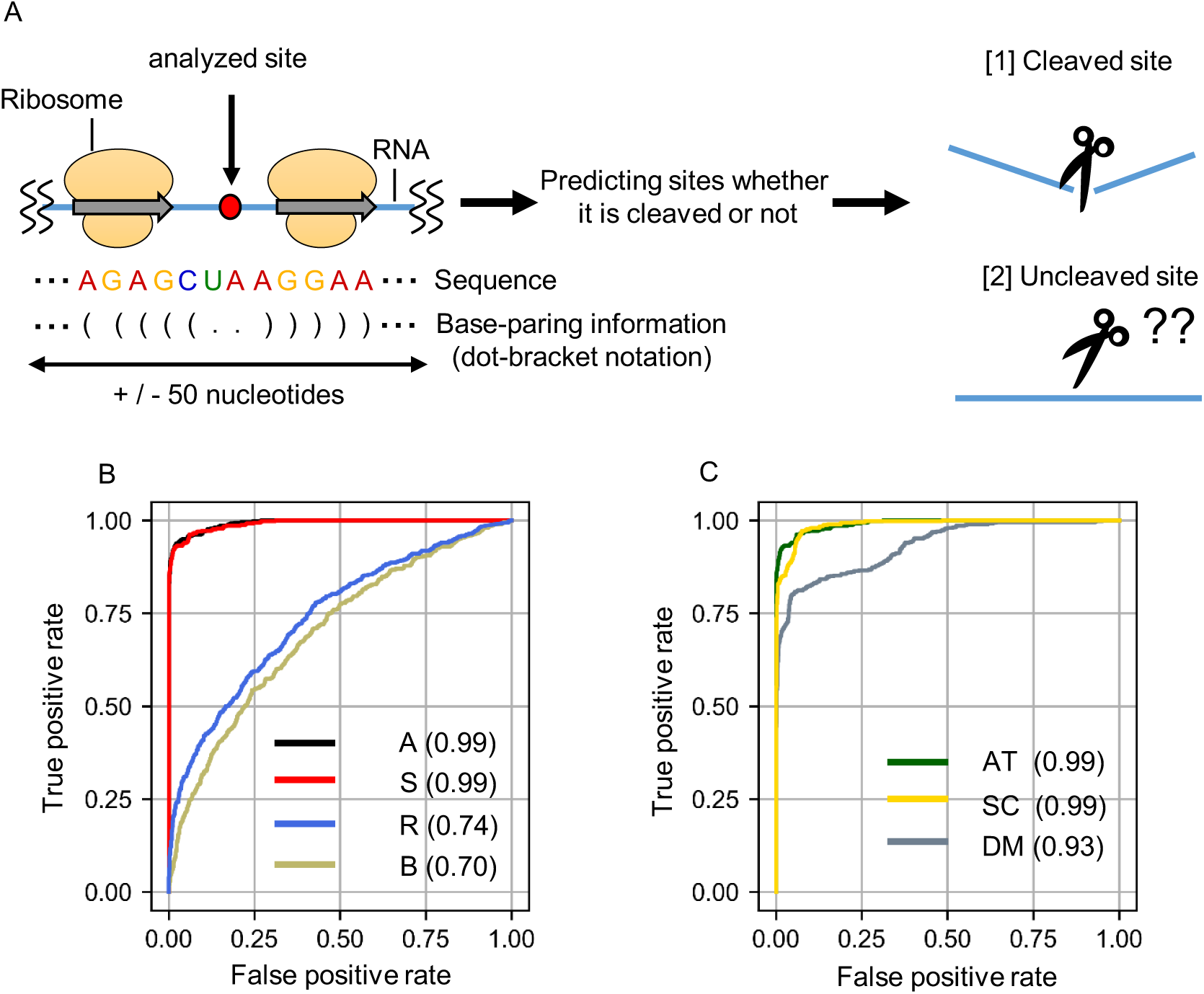
Feature selection using RF classifier. Using information on ribosome occupancy, ribosome position, sequence, and secondary structure (base-pairing information) around cleavage sites, we predicted whether each site was cleaved (A). Data were divided into training and test data sets. We constructed models using the training data and evaluated the accuracy using the test data. Four models were constructed: [A] all features (sequence, ribosome position and ribosome occupancy, base-pairing), [S] only sequence, [R] only ribosome position or ribosome occupancy, and [B] only base-pairing information around the cleavage sites. Models were evaluated using ROC curves and AUC values (B). Parentheses indicate AUC values in each model. We predicted cleaved sites in *D. melanogaster* and *S. cerevisiae* using a model constructed exclusively from *A. thaliana* sequence (C). ROC curves and AUC values in *A. thaliana* are the same as in Figure 6B. AT, *A. thaliana*; SC, *S. cerevisiae*; DM, *D. melanogaster*.

We hypothesized that prediction accuracy would still be high in *D. melanogaster* and *S. cerevisiae* using a model that was constructed in *A. thaliana* using only sequence information, so long as similar sequence features were important for determining cleavage position across eukaryotes. As shown in Figure 6C, AUC values were > 0.9 in *D. melanogaster* and *S. cerevisiae*. Consistent with the sequence motif analysis, prediction accuracy using models constructed in *A. thaliana* were higher in *S. cerevisiae* than in *D. melanogaster*. These results suggest that common sequence features involved in determining cleavage position were highly conserved among eukaryotes, but that species-specific sequences also influence RNA cleavage.

## DISCUSSION

### A High G nucleotide frequencies is a strong indicator for cleavage sites

In TREseq, we observed high G-nucleotide frequency around cleavage sites. Although these sequence features around cleavage sites were not clearly observed in previous studies, weak trends were reported in previous degradome analyses. In Akron-seq, higher G-nucleotide frequency (defined as G-quadruplex) was observed downstream of the cleavage site in mammals, and similar sequence features were also identified by PAREseq or 5PSeq (Ibrahim et al. 2018). In particular, Akron-seq observed the expected G-quadruplex structures in highly conserved regions in vertebrates.

Previous degradome sequencing methods involved ligation of single-stranded adapters using T4 RNA ligase, making it difficult to detect 5’-truncated RNAs with extensive base-pairing (Zhuang et al. 2012). This result is consistent with a previous degradome analysis in which open structures were observed around cleavage sites (Ibrahim et al. 2018). By contrast, TREseq (CAGE) methods use ligation of different adapters, and open structures were not observed around the cleavage sites (Supplementary Figure S13). Therefore, G-rich sequences were observed around cleavage sites, but not downstream of them. Also, because poly (A) RNA selection caused the detected reads or sites to be enriched in 3’ UTRs or around stop codons (Weinberg et al. 2016), evaluation of cleavage efficiencies differed from previous degradome analysis. Because of these improvements in the experimental procedures, specific sequence features around cleavage sites were clearly observed in TREseq. Although we cannot exclude the possibility that G-quadruplex structures affect cleavage efficiencies, RF classifier analyses indicated that the influence of RNA structure was weak (Figure 6). Considering that cleavage sites with higher G-nucleotide frequencies were detected by different methods in different species, we believe that these sequence features are actually present around the cleavage sites, thus indicting that sequence-specific RNA degradation mechanisms are highly conserved in eukaryotes.

Although we focused on uncapped cleavage sites in this study, some cleavage sites are secondarily capped in the cytosol (Schoenberg and Maquat 2009). Indeed, a recapping enzyme has been identified in animals, and some cleaved RNAs that originate from no-go decay (NGD) are actually re-capped (Schoenberg and Maquat 2009). However, because only the Cap RNA library was used in CAGE method, the ratio of Cap to Cap-less RNA could not be calculated. By contrast, we made Cap RNA and Cap-less RNA libraries in TREseq, and can roughly estimate the proportion of re-capped RNA from cleavage site data by comparing libraries. We expanded the annotation region of the Cap RNA data following data processing in TREseq, and calculated the expected re-capped RNA in cleavage site data using exon CDS regions from different species (Supplementary Figure S20). Some cleavage sites in *A. thaliana* appeared to be re-capped RNA, suggesting the existence of re-capped cleavage sites in plants. However, the proportion of re-capped cleavage sites was less than 10% in *A. thaliana, D. melanogaster*, and *S. cerevisiae* (Supplementary Figure S20). These results are consistent with a previous study reporting that thousands of cleavage sites in mammals are Cap-less RNA (Malka et al. 2017), suggesting that the majority of cleavage sites detected in genome-wide analysis are uncapped.

### Ribosome positions have little effect on RNA cleavage position

A previous study reported that ribosome movement is important for RNA stability (Presnyak et al. 2015). In particular, slow ribosome movement (ribosome stalling or pausing) causes endonucleolytic cleavage (Doma and Parker 2006; Simms et al. 2017). Although we do not know whether similar trans-acting factors in NGD are related to cleavage sites detected in genome-wide degradome analysis, we can refer to the relationships between ribosome movement and cleavage sites. NGD can be induced by inserting several codons into a reporter gene, and the resultant cleavage sites can be detected in yeast (Simms et al. 2017; Doma and Parker 2006). These results suggest that the translation process is related to cleavage probability.

Initially, these RNA cleavages were thought to occur near sites of ribosome stalling or pausing detected in the NGD analysis (Doma and Parker 2006). However, RNA cleavage sites were widely identified in the gene, including beside the ribosomes based on the analysis in recent study (Simms et al. 2017). These results suggest that ribosome movement is important for RNA cleavage, but not critical for determining cleavage position. In our analyses, we cannot observe a strong relationship between ribosome and cleavage position. Specifically, we observed that distribution patterns between ribosome and cleavage sites have three-nucleotide periodicities, but the phases were generally not matched (Figure 5). These results were observed not only in TREseq analysis but also in another previous study (Ibrahim et al. 2018; Pelechano et al. 2015). Similar results were observed in feature selection using an RF classifier model in *A. thaliana, D. melanogaster,* and *S. cerevisiae*: ribosome movement seemed to be related to cleavage position, but was not a major determinant (Figure 6), suggesting that ribosomes occupancy has a positive effect on cleavage efficiencies, but ribosome position does not have a strong effect on cleavage position. Because codon frequencies were generally the same in each translation frame in the transcripts, we could observe the three-nucleotide periodicity of nucleotide frequencies in CDS regions in *A. thaliana, D. melanogaster,* and *S. cerevisiae* (Supplementary Figure S21 and S22) and these periodicities appear to form the distribution patterns of cleavage sites.

Previous study reported that these periodicities in yeast and plant are caused by exonucleolytic digestion because the amplitude of periodicity was decreased in exonuclease mutant (Pelechano et al. 2015; Yu et al. 2016). However, even in the exonuclease mutant, the periodicities were still observed in animal (Ibrahim et al. 2018). In addition, disappearance of periodicity appeared to be caused by low coverage of detected 5’ degradation intermediates and our previous study revealed that periodicity of 5’ degradation intermediates was almost identical in W T and exonuclease mutants in *A. thaliana* (Ueno et al. 2020). Also, we obtained mapping data of 5Pseq in yeast (Pelechano et al. 2015) and reanalyzed whether the periodicity disappeared around start or stop codon in exonuclease mutant (Supplementary Figure S23). Because number of detected reads (read count) at each 5’ degradation intermediates were sometimes dependent on its RNA accumulation levels, there was a possibility that the distribution pattern based on the read count did not reflect the genome-wide tendency due to highly expressed genes. Therefore, we calculated the number of 5’ degradation intermediates (detected sites in 5Pseq) at each RNA position by count and frequency (relative to all detected 5’ degradation intermediates). The amplitude of periodicity was slightly decreased around stop codon in exonuclease mutant (Supplementary Figure S23B and D), but the periodicity around start codon was almost similar or more clearly observed in exonuclease mutant compared to W T (Supplementary Figure S23A and C). These results suggest that exonuclease is not major determinant for forming three-nucleotide periodicity in yeast. Considering that periodicity was still observed in exonuclease mutant in plant (Ueno et al. 2020), animal (Ibrahim et al. 2018) and even in yeast (Supplementary Figure S23), periodicity appear to be originated from endonucleolytic cleavage–dependent RNA degradation.

In summary, using TREseq, we observed that cleavage efficiencies play important roles in RNA stability in various species. We observed that sequence features and ribosome occupancy h ad a positive effect on cleavage efficiencies, and that sequence features were more important than ribosome movement for determining RNA cleavage positions across eukaryotes. These results suggest that similar sequence-specific endonucleolytic cleavage mechanisms are conserved across species, and that these mechanisms contribute to the control of RNA stability among eukaryotes. On the other hand, given that different sequences also seemed to be related to RNA cleavage, based on the results of the sequence motifs and RF classifier analyses, further studies are needed to elucidate endonucleolytic cleavage–dependent RNA degradation in eukaryotes.

## MATERIAL AND METHODS

### Growth conditions for *A. thaliana*

*A. thaliana* T87 cell suspension was cultured in modified Murashige–Skoog medium as described previously (Matsui et al. 2011).

### Growth conditions for *D. melanogaster*

S2-R+ cells (Drosophila Genomics Resource Center # 150) were grown in Schneider’s medium supplemented with 10% FBS and 100 U/ ml penicillin–streptomycin at 25°C.

### Growth conditions for *S. cerevisiae*

*S. cerevisiae* strain Σ1278 b was cultured at 30°C to exponential growth phase (OD_600_ of 1.0) in YPD medium containing 2% (wt/vol) glucose, 1% (wt/vol) Difco Bacto yeast extract (Thermo Fisher Scientific, Waltham, MA, USA), and 2% (wt/vol) Difco Bacto peptone (Thermo Fisher Scientific).

### Library construction for truncated RNA-end sequencing (TREseq)

TREseq libraries were constructed using RNA from *D. melanogaster* and *S. cerevisiae* as described previously (Ueno et al. 2018, 2020). In brief, RNA was isolated and purified, and r RNA was depleted using the Ribo-Zero r RNA Removal Kit (Human/ Mouse/ Rat) (I l lumina, San Diego, CA, USA). After r RNA depletion, total RNA was reverse-transcribed with a random primer, and RNA–cDNA hybrids were separated into Cap RNA–cDNA hybrid (Cap RNA) and Cap-less RNA– cDNA hybrid (Cap-less RNA) fractions using the Cap-trapping method. Subsequently, we used different 5’ adapter sequences in Cap and Cap-less RNA (Table 1), followed by library preparation for n An T-i CAGE (Murata et al. 2014; Adiconis et al. 2018). We used Cap-less RNA to detect 5’ degradation intermediates and Cap RNA to estimate transcription start site (TSS) and RNA abundance using more than 50 reads for each gene. The libraries were sequenced on a Next Seq 500 instrument (Illumina).

**Table. 1.**
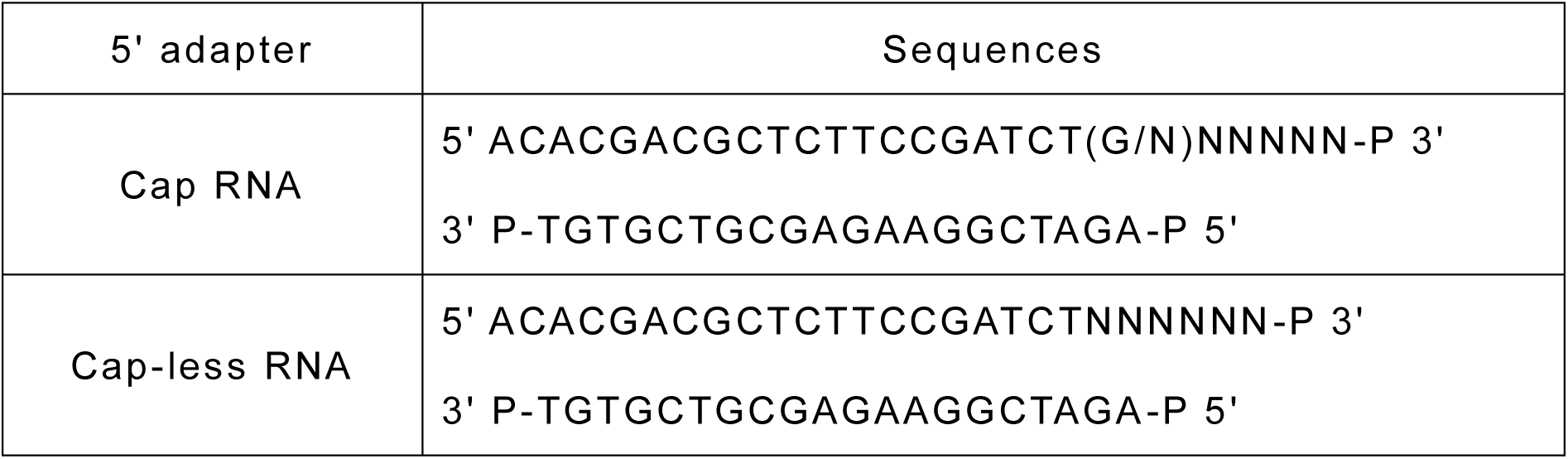
5’ adapter sequence used in TREseq.

For the TREseq library without Cap-trapping method in *A. thaliana,* we did not conduct Cap-trapping method in library preparation and used 5’ adapter sequence similar to Cap-less RNA library (Table 1). The libraries were subjected to sequencing on a Next Seq 500 instrument (Illumina).

### Data processing for TREseq

We used a modified MOIRAI system to process TREseq data (Hasegawa et al. 2014; Ueno et al. 2018, 2020). For calculation of 5’ degradation intermediates (Cap-less RNA), low-quality reads or reads originating from rRNA were removed. The remaining reads were mapped to the TAIR version 10 reference genome (www.arabidopsis.org), Flybase version 6.31 reference genome (https://flybase.org/), or Sigma 1278b_MIT_2009_ACVY01000000 reference genome (https://www.yeastgenome.org/) using HISAT2. After converting BAM files to SAM files, we removed reads with mismatches within three nucleotides of the 5’ end of a mapped read, except for TREseq library without Cap-trapping method. Because the 5’ cap (7-methylguanosine) is not encoded by the genome, most Cap RNAs in the cleavage sites data were removed. Subsequently, the SAM files were converted to BED files, which were used to count only the first nucleotide (5’ end) of each read. Because we used different mapping software than in previous TREseq analysis, data processing for *A. thaliana* is shown in the figures (Supplementary Figure S1). For calculation of TSS or RNA abundance (Cap RNA), the data were processed by CAGE (Murata et al. 2014; Adiconis et al. 2018). In the library preparation step, we could not completely exclude contamination (Cap RNA in Cap-less RNA library or Cap-less RNA in Cap RNA library). Hence, we removed Cap RNA from Cap-less RNA data and vice versa by comparing each library, as described in previous study (Ueno et al. 2020) (Supplementary Figures S6-S8). De-capped RNA was removed from Cap-less RNA data by comparing information for Cap and Cap-less RNA. As an indicator of cleavage efficiencies at each site, we defined the reads at each cleavage site normalized by RNA abundance as the cleavage score at the site level (CS_site_), as described in previous studies (Ueno et al. 2020, 2018). For the gene level, we defined the total CS_site_ value at each gene or each CDS region as the CS_gene_ or CS_CDS_ value, respectively.

### Library construction for ribosome profiling

We selected ribosome-protected fragments (RPFs) as previously described (Lei et al. 2015; Yamasaki et al. 2015). In brief, *A. thaliana* T87 cells were harvested 3 days after inoculation and frozen in liquid nitrogen, followed by homogenization with extraction buffer (200 mM Tris-HCl, p H8.5, 50 mM KCl, 25 mM MgCl_2_, 2 mM EGTA, 100 µg/ml heparin, 100 µg/ml cycloheximide, 2% polyoxyethylene 10-tridecyl ether, 1% sodium deoxycholate), and centrifugation at 15,000 × g for 10 min at 4°C (Lei et al. 2015). We added 6 μl of RNase I (Thermo Fisher Scientific) for 30 min, and stopped the reaction by adding 10 μl of RNase inhibitor (Thermo Fisher Scientific). Using a 26.25–71.25% sucrose density gradient buffer (200 mM Tris-HCl, pH 8.5, 200 mM KCl, 200 mM MgCl_2_), monosomes were collected by sucrose density gradient centrifugation at 55,000 r.p.m. for 50 min at 4°C in an SW55 rotor (Beckman Coulter, Brea, CA, USA). After isolation of monosomes, RPFs were purified using the TruSeq Ribo Profile kit (Illumina). The libraries were sequenced on a Next Seq 500.

### Data processing for ribosome profiling

Adapter sequences were trimmed (Luo et al. 2018; Gerashchenko and Gladyshev 2017) and reads were mapped using the modified MOIRAI system (Ueno et al. 2018, 2020). The remaining reads were mapped to the TAIR version 10 reference genome (www.arabidopsis.org), Flybase version 6.31 reference genome (https://flybase.org/), or Sigma 1278 b_MIT_2009_ACVY01000000 reference genome (https://www.yeastgenome.org/) as appropriate, using BWA. After mapping, we counted the first nucleotide (5’ end) of each read using the BED files. As an indicator of RPFs at each site, we defined the average of reads at each 5’ RPF, normalized against RNA abundance, as ribosome occupancy at the site level (RO_site_). For the gene level, we defined the total RO_site_ value in the CDS region at each gene as the RO_CDS_ value.

### Library construction for Nanopore sequencing

Total RNA was extracted using Trizol (Thermo Fisher Scientific), and mRNA was isolated from total RNA using the Magnosphere Ultra Pure mRNA Purification Kit (Takara Bio, Kusatsu, Shiga, Japan). Nanopore lib raries were prepared from poly(A) ^+^ RNA using the Nanopore PCR cDNA Sequencing Kit (Oxford Nanopore Technologies, Oxford, UK). We conducted 12 cycles of PCR amplification. RNA sequencing on the GridION platform (Oxford Nanopore Technologies) was performed for 48 h on ONT R9.4 flow cells using a minknow-core-gridion 3.1.8 software (Oxford Nanopore Technologies).

### Data processing for Nanopore profiling

Reads were base-called with ont-guppy version 2.0.8-1 (Oxford Nanopore Technologies). Fully sequenced reads were identified and oriented using Pychopper2 (Oxford Nanopore Technologies). Reads were mapped to the Arabidopsis TAIR10 genome using Minimap2. Subsequently, the SAM files were converted to BED files, which were used to count only the first nucleotide (5’ end) of each read.

### *In vitro* synthesized RNA and quantitative RT-PCR (qRT-PCR) analysis

*In vitro* synthesis of uncapped *Renilla* luciferase (*R-luc)* from plasmid pT3-RL-pA was performed as described previously (Matsuura et al. 2008). We added 1 ng of uncapped *R-luc* RNA to 5 μl of total RNA and conducted rRNA depletion and reverse transcription as described for Cap-less RNA library preparation in TREseq. We conducted qRT-PCR on a LightCycler 480 (Roche Applied Science, Upper Bavaria, Germany). Gene-specific primers used for qRT– PCR of *R-luc* RNA were 5’-GGATTCTTTTCCAATGCTATTGTT-3’ and 5’-AAGACCTTTTACTTTGACAAATTCAGT-3’ (Matsuura et al. 2008).

### Data analysis

For motif analysis, sequence motifs adjacent to the cleavage sites were identified using MEME motif analysis tools (http://meme-suite.org). Motif distribution was calculated using FIMO.

As for GO enrichment analysis, we used the Gene Ontology enRIchment anaLysis and visuaLizAtion tool (GORILLA) (http://cbl-gorilla.cs.technion.ac.il/).

### Construction of the random forest (RF) classifier

For feature selection, we used the RandomForestClassifier from the Python package scikit-learn. We defined the highest CS_site_ value in each gene as the cleavage site and selected a total of 1000 genes. For non-cleaved sites, we selected sites from the selected 1000 genes with no CS_site_ value at least around +/- 30 nucleotide. We divided data into training (500 genes) and test data (500 genes) (Table 2). In the models, we used three features: [1] sequence, [2] secondary structure, and [3] ribosome position and ribosome occupancy. We converted sequences or secondary structure information into numeric numbers for the RF classifier. We optimized parameters of the RF classifier based on the out-of-the-bag (OOB) score in the training data and receiver operating characteristic (ROC) curves and area under the curve (AUC) values in the test data, as described in previous studies (Lloréns-Rico et al. 2015; Singh et al. 2017). The ROC curve specifies the true positive rate and false-positive rate; a high AUC value indicates high prediction accuracy. We constructed an RF classifier using the training data and predicted accuracy using the test data. We also predicted cleavage sites in *D. melanogaster* and *S. cerevisiae* using the RF classifier constructed in *A. thaliana*.

**Table 2.** Sites analyzed in RF classifier model

|  | *A. thaliana* |  | *D. melanogaster* |  | *S. cerevisiae* |  |
| --- | --- | --- | --- | --- | --- | --- |
|  | Training | Test | Training | Test | Training | Test |
| Cleaved sites | 500 | 500 | 500 | 500 | 500 | 500 |
| Uncleaved sites | 1926 | 1732 | 3675 | 3327 | 2321 | 2314 |

## DATA ACCESS

TREseq reads are available in the DDBJ Sequence Read Archive (DRA) database with accession numbers DRA 010805 (*Drosophila melanogaster*), DRA 010806 (*Saccharomyces cerevisiae*) and DRA 010923 (*Arabidopsis thaliana* without Cap-trapping method). Ribosome profiling and Nanopore sequencing reads in *A. thaliana* are available with accession numbers DRA 010802 and DRA 010803, respectively.

## COMPETING INTEREST STATEMENT

The authors declare no competing interests.

## ACKNOWLEDGMENTS

We thank the Okamura and Takagi laboratories at the Nara Institute of Science and Technology (NAIST) for helpful technical advice regarding S2-R+ cells and strain Σ1278b. We also thank DNAFORM for excellent deep sequencing analysis and helpful suggestions, and Yoichiro Watanabe at the NAIST for useful discussion and proofreading.

## Notes

### Competing Interest Statement

The authors have declared no competing interest.

